# The TARGET OF RAPAMYCIN (TOR) kinase controls shade-mediated hypocotyl elongation responses in Arabidopsis

**DOI:** 10.64898/2026.06.26.734770

**Authors:** Sarah Courbier, Mikhail Schepetilnikov, Sebastian N. W. Hoernstein, Isabel Lembke, Christian Meyer, Pitter F. Huesgen, Andreas Hiltbrunner

**Affiliations:** Faculty of Biology, University of Freiburg, 79104, Freiburg, Germany; CIBSS - Center for Integrative Biological Signalling Studies, University of Freiburg, 79104, Freiburg, Germany; Institut de biologie moléculaire des plantes UPR2357 (CNRS) Université de Strasbourg, France; Institute Jean-Pierre Bourgin for Plant Sciences (IJPB), INRAe, AgroParisTech, Université Paris-Saclay, Versailles, France

**Author notes:** Senior author (Andreas Hiltbrunner).

**Keywords:** TOR, shade avoidance, growth, plasticity, translation

## Abstract

Plants perceive neighboring vegetation through an enrichment of far-red light (shade) in the environment. These changes in light quality trigger molecular and physiological responses aimed at outgrowing competitors, collectively known as the shade avoidance syndrome. In this study, we identify the TARGET OF RAPAMYCIN (TOR) complex 1 (TORC1), a major growth-regulating hub in eukaryotes, as a driver of shade-mediated growth responses in plants. Combining physiology, genetics, biochemistry, and proteomics, we show that TOR activity is rapidly enhanced upon shade perception and is required for proper shade responses, as TOR inhibition severely impairs shade-mediated elongation. Furthermore, we found that the control of shade-mediated elongation by TOR involves auxin-dependent mechanisms, requires efficient translation activity, and is closely linked to epidermal cell elongation capacity. Altogether, our work identifies TOR as a key integrator of light quality signals to control adaptive growth responses. Finally, we further highlight the conservation of shade-mediated TOR activation in tomato, with potential implications for engineering crop cultivars better suited to high-density planting.

## Introduction

Light is crucial for plants to navigate through their life cycle. While light availability is essential to fuel photosynthesis, light quality is equally important for plants to assess their surroundings and make adequate adaptive decisions. Light quality is sensed by a sophisticated set of photoreceptors ranging from the UV-B receptor UVR8, blue light photoreceptors phototropins and cryptochromes, to red and far-red light photoreceptors phytochromes. Phytochromes are synthesized in their inactive Pr form in the cytosol and, upon light-induced conversion to the active Pfr form, translocate to the nucleus to trigger downstream molecular and physiological responses. In *Arabidopsis thaliana* (Arabidopsis), five phytochromes (phyA-phyE) coordinate responses to red and far-red light, with phyB playing a major role in neighbor proximity responses ^1^.

At high planting density, plants absorb blue and red light while they reflect or transmit far-red light, in turn creating a far-red enriched light environment. As a result, the ratio between red and far-red light drops (low R:FR) and triggers adaptive growth responses in plants known as the shade avoidance syndrome, which is well characterized in Arabidopsis ^2,3^. Under low R:FR conditions, the photoinactivation of phyB releases PHYTOCHROME-INTERACTING FACTORs (PIFs) from sequestration. These transcription factors act as positive regulators of elongation and thereby allow downstream induction of growth-related genes. In Arabidopsis seedlings, low R:FR perception activates rapid auxin biosynthesis, signaling and transport from the cotyledons towards the hypocotyl to trigger elongation growth. More specifically, auxin is transported via PIN-FORMED (PIN) auxin efflux transporters, mainly PIN3, which relocalizes from the periclinal to the anticlinal poles of pericycle cells, thereby transporting auxin radially from the vascular tissue towards the hypocotyl epidermal cells ^4,5^. In adult plants, local far-red light enrichment applied to the leaf tip triggers long distance auxin transport from the tip towards the abaxial side of the petiole to trigger hyponasty ^6,7^. Since the 1970s, auxin is believed to promote elongation growth via the activation of plasma membrane-localized H+-ATPases, leading to H+ efflux through the plasma membrane, apoplastic pH acidification, induction of the cell wall remodeling machinery, and cell wall loosening in both roots and hypocotyls ^8–10^. Epidermal cell elongation also involves brassinosteroid signaling, which by crosstalk with auxin acts as a positive regulator of elongation growth ^5^.

TARGET OF RAPAMYCIN (TOR) is a conserved serine/threonine protein kinase that functions as a central growth regulator in eukaryotes. In plants, TOR forms a single complex, TORC1, with Regulatory-Associated Protein of mTOR (RAPTOR) and Lethal with Sec Thirteen 8 (LST8). TORC1 is essential for plants as loss-of-function *tor* mutants are embryo lethal and plants lacking *RAPTOR* display major post-embryonic growth, hormonal and metabolic defects ^11–13^. Modulation of TOR activity in response to endogenous and exogenous signals is key for plant growth, development and adaptation. On one hand, active TOR promotes cell division, facilitates cell-to-cell transport (through plasmodesmata), and contributes to ribosome biogenesis and translation. On the other hand, active TOR negatively regulates senescence, biotic stress responses and autophagy in coordination with the metabolic sensor Sucrose non-fermentable 1 (SNF1)-related protein kinase 1 (SnRK1) ^14–16^.

While factors such as auxin, light, sugars, nutrients, and components of the circadian clock have been shown to promote the activity of the TOR complex and be linked to development ^13,17–19^, the molecular mechanisms linking upstream signals to TOR activation remain unclear. Still, several candidate regulators have been identified, including the small GTPase ROP2, which links auxin to TOR activation, as auxin fails to activate TOR-mediated translation in the *rop2* mutant background ^20^. The ROP2-TOR axis was also associated with nutrient homeostasis, as nitrogen sources and amino acids (*i.e* glutamine) activate TOR ^21^. Furthermore, TOR inhibition by AZD8055 decreased ammonium uptake by plant roots and resulted in the accumulation of glutamine ^22^. In the shoot, although sugar-activated TOR is sufficient to drive shoot elongation in darkness (skotomorphogenesis) ^23^, TOR activation by light and photosynthesis-derived sugars usually goes hand in hand. Consistent with its central role in coordinating growth with carbon availability, glucose-TOR signaling is required to activate meristem activity and promote the development of new organs ^24^. Sugar-TOR signaling promotes the accumulation of the brassinosteroid (BR)-signaling transcription factor BZR1 by limiting its degradation via autophagy, whereas TOR inhibition is associated with decreased BZR1 abundance, thereby reducing hypocotyl elongation responses ^25^. TORC1 also plays a critical role in plant responses to fluctuating light conditions. LST8 is required for adaptation to long-day photoperiods ^26^, while RAPTOR, particularly RAPTOR1B, is involved in photoperiod sensing and flowering induction as *raptor1b* mutants are unable to initiate the floral transition under long-day light regimes ^27^. Furthermore, variations in light intensity and site of illumination (root, shoot or both) have been shown to regulate plant growth via differential TOR activity ^28^.

Only a few studies have demonstrated a link between light quality and TOR signaling. In Arabidopsis seedlings, monochromatic blue and FR light treatments promote TOR activity compared with darkness in a phyA- and CRY-dependent manner, thereby controlling apical hook unfolding ^29^. More recently, TOR activation by sucrose was shown to alleviate phyA-mediated inhibition of leaf expansion under deep shade conditions (low PAR + low R:FR) ^30^. Previous work in tomato also showed *TOR* expression to be upregulated by low R:FR conditions, thereby raising the possibility that TOR kinase activity contributes to the onset of shade-mediated growth responses ^31^. Despite emerging evidence, our understanding of how light quality regulates TOR dynamics remains limited. In particular, how TOR activity dynamics in response to changes in light quality contribute to adaptive growth responses remains largely unexplored. Shade avoidance is a brilliant example of rapid adaptive growth in plants. Therefore, unraveling whether and how TOR integrates shade signals to trigger physiological and morphological adaptation is of particular interest, especially in the context of agriculture.

In this study, we identify TOR as a key component of shade-mediated hypocotyl elongation growth in *Arabidopsis thaliana* seedlings. We found that FR-enriched light conditions promote TOR activity and that TOR inhibition limits shade-induced hypocotyl elongation growth responses. We conclude that the activation of TOR by FR-enriched conditions promotes global translation efficiency required for rapid growth responses, ultimately affecting auxin homeostasis and cell wall elongation processes. While the regulatory network through which TOR controls adaptive growth responses to shade still requires further clarification, our findings bridge the fields of light and TORC1 signaling by demonstrating their physiological integration in the regulation of shade responses.

## Results

### Far-red light enrichment promotes TOR activity

To assess whether TOR activity is affected by changes in light quality, especially in the context of neighbor proximity, we quantified hypocotyl growth in *Arabidopsis thaliana* Col-0 (Arabidopsis) seedlings grown either in continuous white light for 7 days (cWL) or in cWL for 4 days followed by 3 days in cWL supplemented with far-red light (cWL+FR) to simulate proximity shade (Fig. S1). As a proxy to assess TOR activity under these light conditions, we used phosphorylation of 40S ribosomal protein S6 (RPS6/eS6) at Ser240, a direct downstream target of TOR-S6K1 signaling ^32^. In parallel with shade-mediated hypocotyl elongation (Fig. 1a and b), we observed a rapid increase in TOR activity after 6 hours of shade treatment, which persisted for at least 24 hours (Fig. 1c), suggesting a link between shade-mediated growth responses and TOR activation. Similarly, seedlings lacking phytochrome B (*phyB-9*), which display a constitutive shade avoidance phenotype, also showed increased TOR activity even in the absence of shade (Fig. S2a-c), further suggesting a role of TOR activity dynamics in shade-mediated elongation responses downstream of phyB inactivation. In adult plants, a connection between increased TOR activity and hyponasty or petiole elongation growth may exist as TOR activity slightly increased in response to WL+FR in leaf tissue compared with WL controls (Fig. S2d-g). Altogether, these results show that FR-enriched conditions promote TOR activity in the hypocotyls of Arabidopsis seedlings, correlating with the activation of shade-mediated growth responses and potentially positioning TOR as a key integrator of environmental signals.

**Figure 1:**
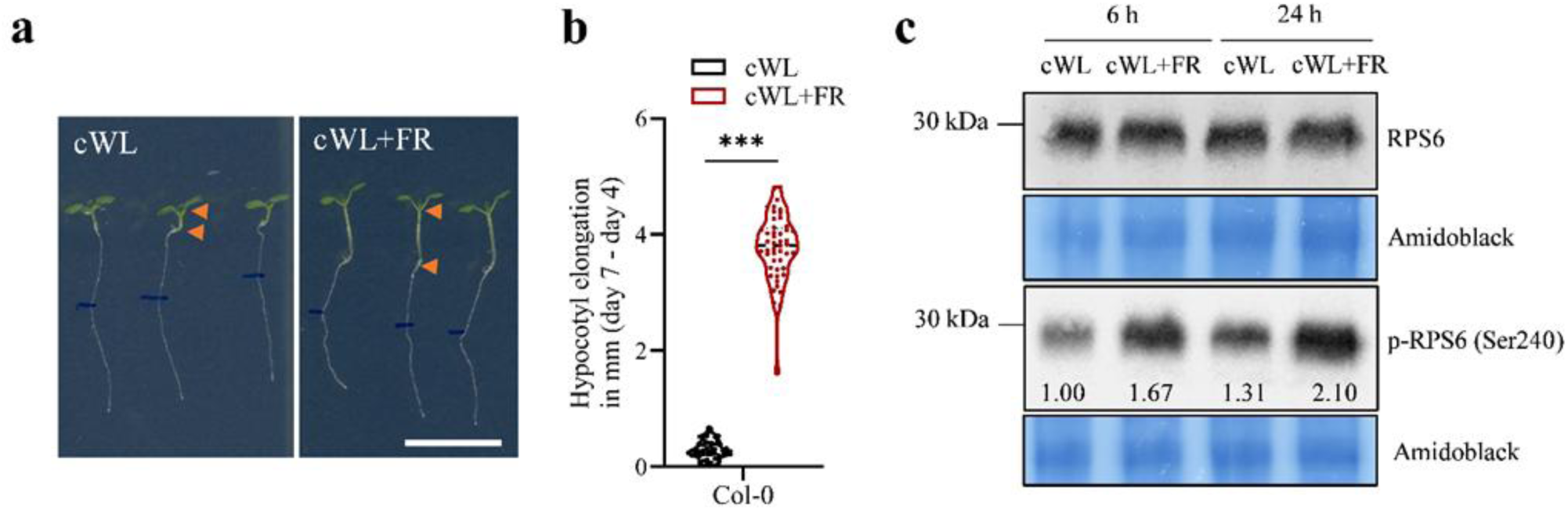
Far-red enrichment stimulates TOR activity. Representative images **(a)** and hypocotyl elongation measurement **(b)** of 7-day-old Col-0 wild-type seedlings after 7 days in continuous white light (cWL; left panel), or after 4 days of cWL followed by 3 days in cWL supplemented with far-red light (cWL+FR; right panel) to simulate canopy shading. Data represent the hypocotyl elongation between before and after 3 days of cWL or cWL+FR treatment. Scale bar = 1 cm. Orange arrows delimit the hypocotyl region. *** corresponds to *p value* < 0.001 according to Student’s t-test. n = 48 hypocotyls measured per conditions. **(c)** Immunoblotting using antibodies against the TOR target 40S ribosomal protein S6 (RPS6A) in its unphosphorylated (anti-RPS6) and phosphorylated (anti-p-RPS6 (Ser240)) state from crude protein extracts (10 μg) originating from 7-day-old Col-0 hypocotyls after 7 days in cWL or 4 days in cWL + 3 days in cWL+FR. Sampling was performed 6 h and 24 h after the start of the shade/AZD8055 treatment. Amidoblack staining was used as a loading control. n = 30-36 hypocotyls per timepoints and conditions. Quantifications are indicated by the ratio in band intensity for p-RPS6 compared with RPS6 for each condition.

### TOR inhibition limits shade-mediated hypocotyl elongation growth

As TOR activity is promoted by FR-enriched light conditions (Fig. 1c), we hypothesized that TOR activation by cWL+FR is required for the proper onset of shade-mediated elongation growth. To test this, we used a pharmacological approach by growing Arabidopsis Col-0 seedlings in cWL for 4 days, prior to transferring them onto new plates containing either DMSO as a control or the specific TOR inhibitor AZD8055 at concentrations of 0.5, 1, 2 or 5 μM. The plates were then incubated in cWL or transferred to cWL+FR (Fig.2; Fig S3). We observed that TOR inhibition had no effect on hypocotyl growth in cWL (Fig. 2a and 2b). In contrast, increasing concentrations of AZD8055 specifically correlated with a progressive reduction of shade-mediated hypocotyl elongation, suggesting that TOR is required for proper elongation responses to shade. In addition, we also observed a typical main root growth inhibition upon shade exposure in mock conditions (Fig. 2c) ^33^, and increasing AZD8055 concentrations affected root elongation growth even further with no difference between control and shaded plants (Fig. 2c). Next, we focused on 1 µM AZD8055, as this concentration decreases elongation by half, therefore allowing us to witness either a rescue or a further decrease in growth phenotypes, while TOR activity remained strongly inhibited under these conditions (Fig. 2d). Consistent with the shorter hypocotyl phenotypes observed upon TOR inhibition, the *rap78* mutant, deficient in TORC1 component <u>R</u>egulatory-<u>A</u>ssociated <u>P</u>rotein of <u>T</u>arget <u>o</u>f <u>R</u>apamycin (*RAPTOR 1B*), and the estradiol-inducible TOR RNAi line *tor-es1* ^34^ displayed reduced shade-mediated hypocotyl elongation compared with wild-type or non-elicited plants (Fig. 2e and 2f). The elongation in response to cWL and cWL+FR were identical between Col-0 and non-elicited *tor-es1*, and both lines were sensitive to AZD8055 to a similar degree (Fig. 2f). In addition, hypocotyl elongation in shade-treated wild-type and cWL-grown *phyB-9* seedlings was similarly reduced by TOR inhibition, further supporting a role for TOR in phyB-controlled growth responses (Fig. 2g). Similarly, elevated temperature also promotes phyB inactivation and stimulates hypocotyl elongation ^35^. Interestingly, TOR also appears to contribute to temperature-mediated elongation response, as AZD8055 similarly repressed both shade- and temperature-induced hypocotyl elongation (Fig. 2h). Altogether, our results suggest a general role for TOR in phyB-controlled regulation of hypocotyl growth.

**Figure 2:**
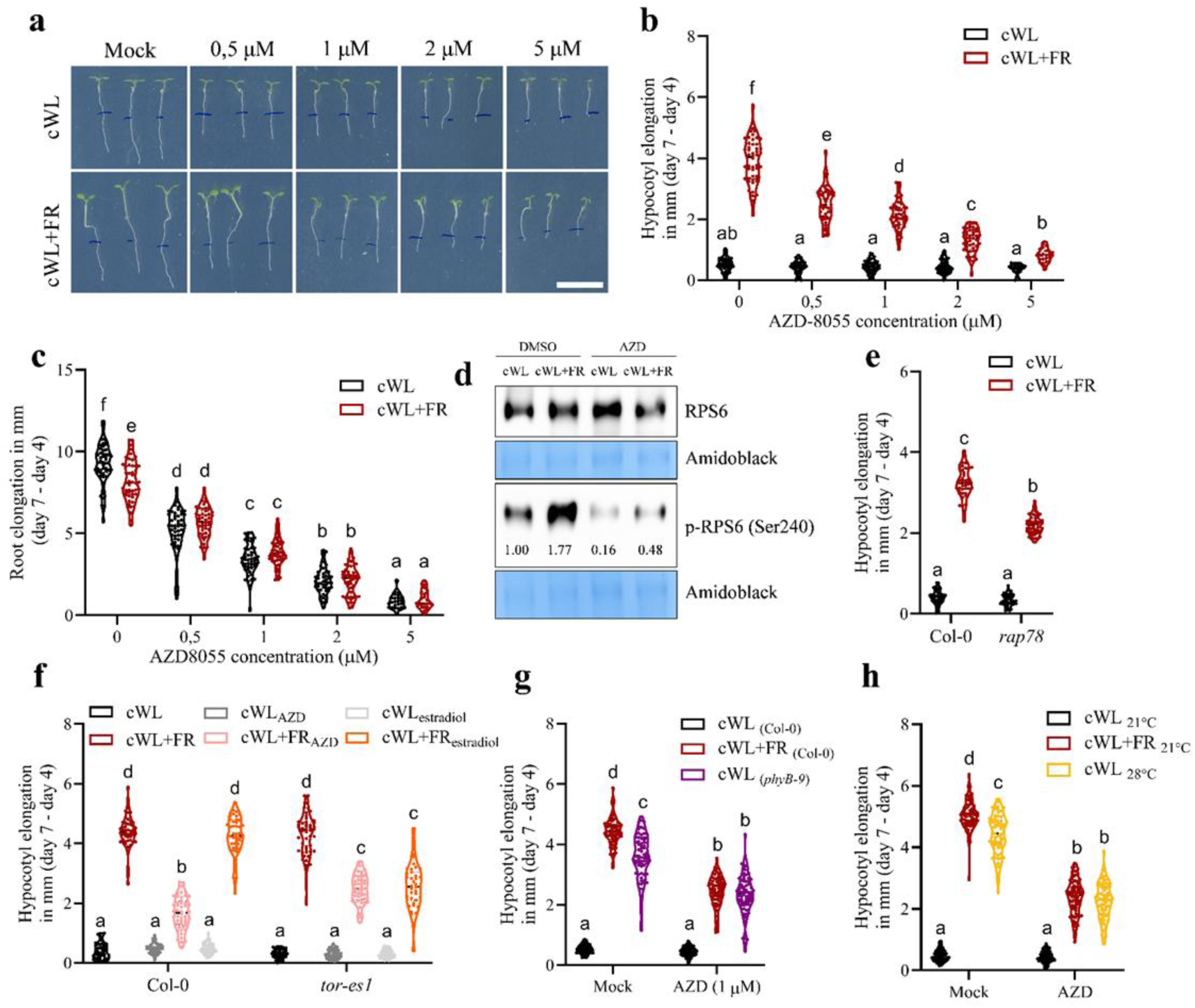
TOR inhibition limits shade-mediated hypocotyl elongation growth. **(a-c) (a)** Representative images, **(b)** hypocotyl elongation **(c)** and root elongation measurement of 7-day-old Col-0 wild-type seedlings after 7 days in continuous white light (cWL), or after 4 days of cWL followed by 3 days in cWL supplemented with far-red light (cWL+FR), and treated with DMSO as a control (0) or with 0.5, 1, 2 and 5 µM of AZD8055. Data represent the elongation between before and after 3 days of cWL or cWL+FR treatment (day 7 – day 4). Root tips were marked immediately after transfer at day 4 and subsequent root elongation was measured. Scale bar = 1 cm. **(d)** Immunoblotting using antibodies against the TOR target 40S ribosomal protein S6 A (RPS6) or against RPS6A phosphorylated on Serine 240 (p-RPS6 (Ser240)). 25 µg of crude protein extracts originating from hypocotyls of Col-0 grown for 7 days in cWL or 4 days in cWL + 3 days in cWL+FR in the presence or absence of 1µM of AZD8055. Sampling was performed 24 h after the start of the shade treatment. Amidoblack staining was used as a loading control. n = 30-36 hypocotyls per conditions. Quantifications are indicated by the ratio in signal intensity for p-RPS6 compared with RPS6 for each condition and normalized to cWL control. **(e-h)** Measurements of hypocotyl length for **(e)** the *RAPTOR1 B* mutant *rap78* grown in cWL or cWL+FR, **(f)** the estradiol-inducible RNAi line *tor-es1* grown in cWL or cWL+FR in the presence and absence of AZD8055 (1 μM) or estradiol (10 μM), **(g)** Col-0 and *phyB-9* mutants in the presence or absence of AZD8055 (1 μM) and **(h)** Col-0 seedlings grown in cWL to either 21 or 28 degrees in the presence and absence of AZD8055 (1 μM). Data represent hypocotyl elongation between day 4 and day 7, *i.e.* before and after 3 days of cWL or cWL+FR treatment. DMSO was used as a control. Scale bar = 1 cm. Letters indicate significant differences according to two-way **(b, c, e)** and three-way **(f, g, h)** ANOVA, Tukey’s post-hoc test. n = 48 hypocotyls measured per conditions.

TOR-dependent growth was recently shown to occur in bryophytes such as *Physcomitrium patens*, suggesting a strong degree of conservation throughout green lineages^36^. In order to test whether TOR activation by shade could be conserved among other shade avoidant plant species with high agronomical impact, we used 3-week-old tomato plants *c.v.* Moneymaker grown in WL (under a long day light regime) and transferred to either WL or WL+FR for 3 days. Under these conditions, plants nicely displayed established shade avoidance traits such as petiole, internode and stem elongation in response to shade (Fig. S4a). Interestingly, we also observed an increase in TOR activity after 6 hours in shade, which is dampened upon AZD8055 treatment (Fig. S4b).

### TOR inhibition limits hypocotyl epidermal cell elongation in response to shade

To investigate how TOR controls shade-mediated elongation responses, we assessed whether TOR inhibition impairs hypocotyl elongation by limiting epidermal cell elongation capacity. Since shade-mediated hypocotyl growth depends on rapid cell elongation rather than cell division, we measured hypocotyl epidermal cell length under cWL or cWL+FR, in the presence or absence of the TOR inhibitor AZD8055 (Fig. 3). As expected, seedlings exposed to cWL+FR displayed longer epidermal cells compared with cWL-grown seedlings. TOR inhibition had no effect on epidermal cell length in cWL, while it drastically reduced cell elongation capacity in cWL+FR (Fig. 3a-c). In addition to affecting epidermal cell elongation, TOR inhibition also appeared to limit hypocotyl radial expansion (Fig. 3d).

**Figure 3:**
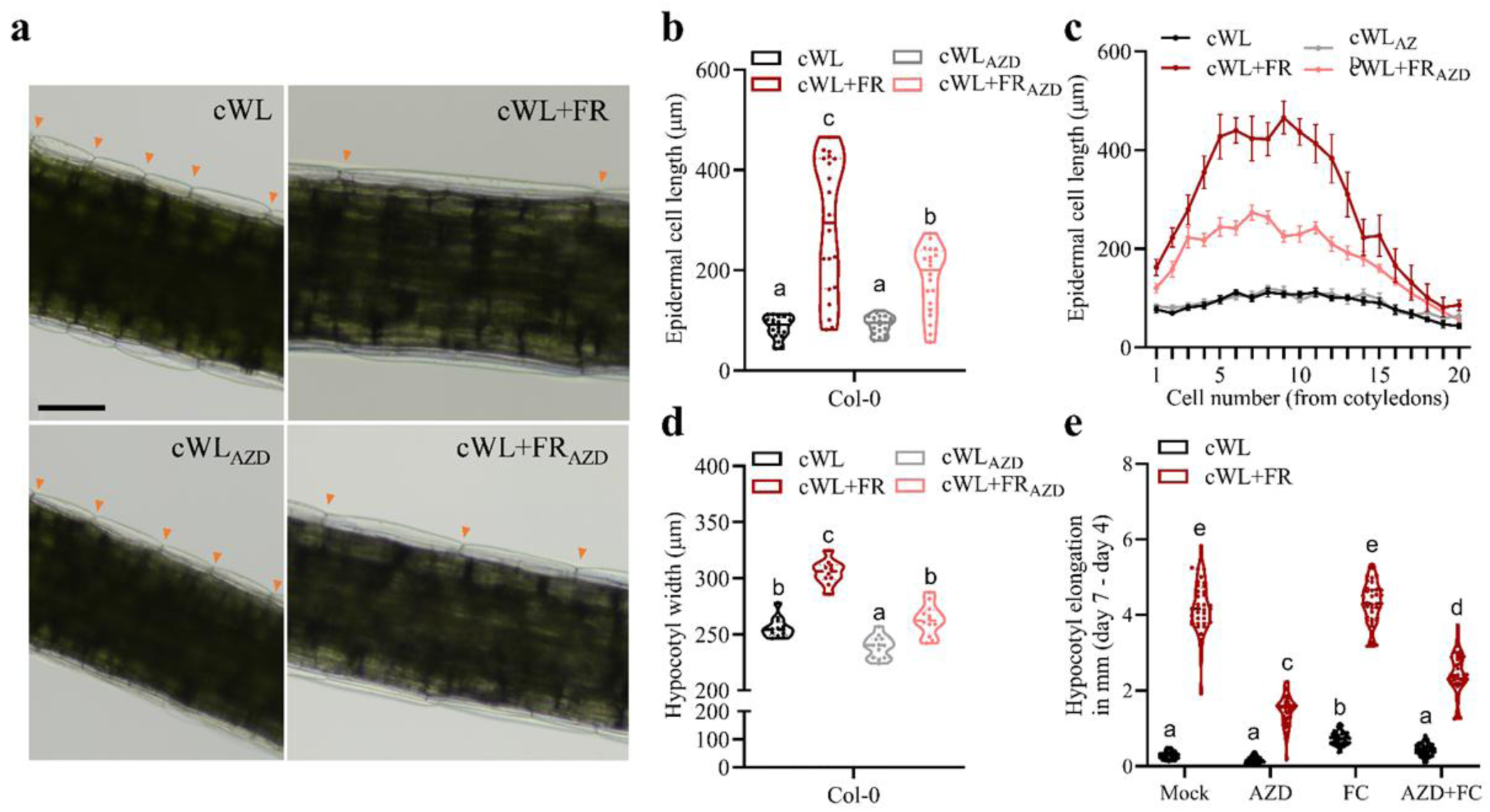
TOR inhibition limits hypocotyl epidermal cell elongation in response to shade. **(a)** Close up light microscopy images of hypocotyls epidermal cells, **(b)** average and **(c)** individual cell length measurements across the hypocotyl and **(d)** hypocotyl width measurements. **(e)** Col-0 hypocotyl elongation measurements upon treatment with AZD-8055 or/and the plasma membrane H^+^-ATPase activator Fusicoccin (FC). All measurements were performed on 7-day-old Col-0 wild-type seedlings grown in continuous white light (cWL) or 4 days of cWL followed by 3 days in cWL supplemented with far-red light (cWL+FR) in the presence and absence of 1 μM AZD8055 (AZD). Letters indicate significant differences according to two-way **(b, d)** or three-way **(e)** ANOVA, Tukey’s post-hoc test. n=10-20 hypocotyls measured per conditions. Orange arrows indicate epidermal cell borders. Scale bar= 200 μm.

Hypocotyl elongation is commonly attributed to the “acid growth theory”, a concept first proposed in the 1970s ^8^ and continuously corroborated and challenged ever since. The model is based on the activation of plasma membrane-localized H+-ATPases by the growth hormone auxin, which triggers H+ efflux through the plasma membrane. This leads to apoplastic pH acidification and subsequent cell wall loosening thereby enabling cell expansion and growth ^10^. To test whether the reduced elongation capacity of AZD8055-treated plants could involve impaired acid growth, we supplemented the growth medium with either AZD8055, the plasma membrane H+-ATPase activator fusicoccin to stimulate apoplastic pH acidification, or both. Interestingly, while AZD8055-treated seedlings displayed impaired shade-mediated elongation, the addition of fusicoccin partially restored their ability to elongate in shade despite TOR inhibition (Fig. 3e). In contrast, cWL-treated seedlings did not respond to any of the treatments, except for an increase in elongation following fusicoccin treatment, in line with its positive effect on cell elongation. Altogether, our data suggest a connection, albeit likely indirect, between TOR activity dynamics and the capacity of hypocotyl epidermal cells to elongate in response to shade.

### TOR is not required for proper shade-induced auxin responses in the hypocotyl

As auxin is known to activate the TOR complex ^20,37^ and to be necessary for shade-induced growth responses, we hypothesized that TOR inhibition limits hypocotyl elongation under shade conditions by reducing auxin signaling activation. To test this, we used a *pDR5::GUS* reporter line to visualize auxin-mediated transcriptional responses in seedlings grown in cWL or exposed to cWL+FR, in the presence or absence of AZD8055, for 24 h prior to GUS staining (Fig. 4a and b). We observed a slight yet significant increase in GUS signal in the hypocotyls of cWL-grown seedlings treated with AZD8055 compared with untreated seedlings. As expected in cWL+FR, the signal predominantly accumulated in the cotyledons, newly formed leaves and leaf primordia. Surprisingly, AZD8055 treatment under cWL+FR conditions triggered GUS signal accumulation in and around the vascular tissue of the hypocotyl, which was completely absent in control plants (Fig. 4a and b).

**Figure 4:**
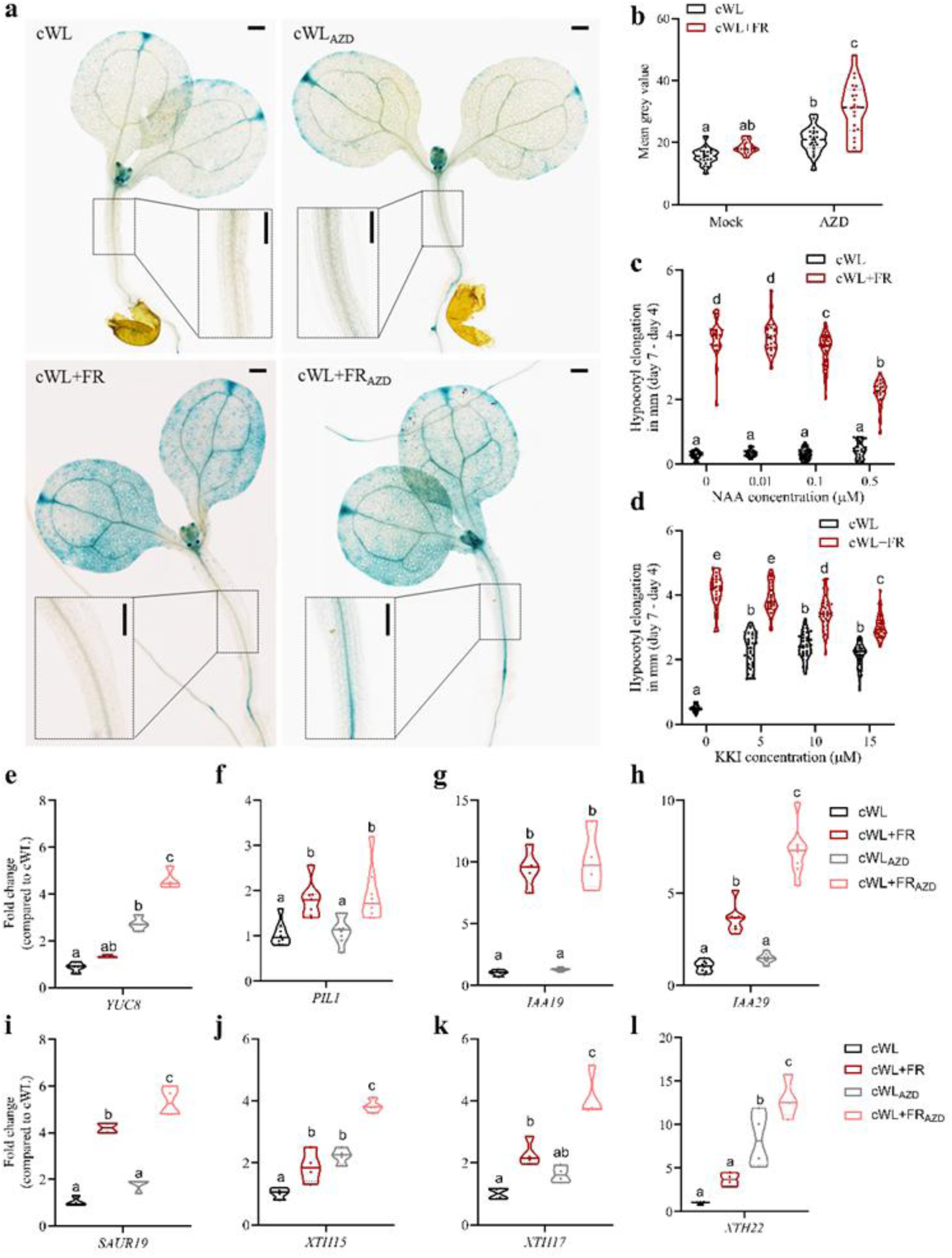
TOR affects auxin signaling and homeostasis in the hypocotyl. **(a and b)** Representative images and quantifications of GUS signal in DR5::GUS seedlings grown for 5 days in continuous white light (cWL) or 4 days in cWL followed by 24 h in cWL supplemented with far-red light (cWL+FR). The GUS signal in the area indicated by dashed boxes was quantified. n = 10 - 20 hypocotyls were measured per conditions. Scale bar = 200 μm. **(c and d)** Hypocotyl elongation in 7-day-old Col-0 wild-type seedlings grown in cWL or 4 days of cWL followed by 3 days in cWL+FR in the presence and absence of **(c)** the synthetic auxin 1-Naphthaleneacetic acid (NAA) or **(d)** the auxin conjugation inhibitor kakeimide (KKI). n > 70 hypocotyls measured per conditions. **(e-l)** Quantification of gene expression in dissected hypocotyls from 7-day old seedlings exposed to cWL or cWL+FR for 24 h, in the presence or absence of the TOR inhibitor AZD8055 (1 µM). Fold-change of expression of genes involved in auxin biosynthesis and signaling as well as genes required for elongation growth was calculated using the 2^-ΔΔCt^ method, taking cWL as reference. Letters indicate significant differences according to two-way ANOVA, Tukey’s post-hoc test.

Since these data did not meet our initial hypothesis, we further tested whether instead of limiting auxin signaling, TOR inhibition could overstimulate auxin signaling, thereby attenuating its growth-promoting effects. Therefore, we grew seedlings on media supplemented with increasing concentrations of the synthetic auxin 1-Naphthaleneacetic acid (NAA), or the auxin conjugation inhibitor kakeimide (KKI), to either promote auxin signaling or increase endogenous free IAA levels, respectively (Fig. 4c and 4d). In particular, KKI promoted hypocotyl elongation under cWL, whereas both chemicals strongly suppressed shade-induced growth in a dose-dependent manner.

To assess whether and how TOR inhibition could affect shade-induced hypocotyl growth responses in an auxin-dependent manner, we quantified transcript levels of genes related to auxin biosynthesis (*YUC8*), auxin signaling and responses (*PIL1/IAA19/IAA29/SAUR19*), and cell expansion (*XTH15,17* and *22*) in hypocotyls of seedlings grown under shade conditions in the presence or absence of TOR inhibition (Fig. 4 e-l). Consistent with the increased GUS signal in the hypocotyls of AZD8055-treated, shade-exposed seedlings, transcript levels of genes involved in auxin biosynthesis, signaling, and cell expansion were upregulated by shade treatment irrespective of TOR inhibition. Interestingly, many of these genes showed the highest expression levels in the hypocotyls of seedlings subjected to combined shade and AZD8055 treatment.

Next, Col-0 seedlings were grown under either cWL or cWL+FR and treated with increasing concentrations of the auxin analog picloram, in the presence or absence of AZD8055 (Fig. S5). Under cWL conditions, increasing picloram concentrations promoted hypocotyl elongation, although this response was less pronounced upon TOR inhibition. However, TOR inhibition restricted elongation in shade conditions, regardless of the picloram concentration applied, suggesting a possible role of TOR in auxin sensitivity (Fig. S5). Altogether, we found that genes involved in auxin biosynthesis, signaling and responses are properly upregulated under shade conditions regardless of TOR activity. However, this transcriptional shade response does not translate into hypocotyl elongation.

### TOR inhibition reshapes the proteome landscape in hypocotyls in response to shade

We next determined how TOR inhibition alters shade-induced changes in the hypocotyl proteome. Col-0 seedlings were grown for 7 days under standard growth conditions (cWL) and then transferred onto fresh plates containing either 1 μM AZD8055 or DMSO as a control. Seedlings were kept in cWL overnight to recover from the transfer and to ensure TOR inhibition before being exposed to cWL+FR conditions or remaining in cWL for 6 h. Hypocotyls were then dissected prior to protein extraction and label-free quantitative proteomic analysis (Fig. 5a).

**Figure 5:**
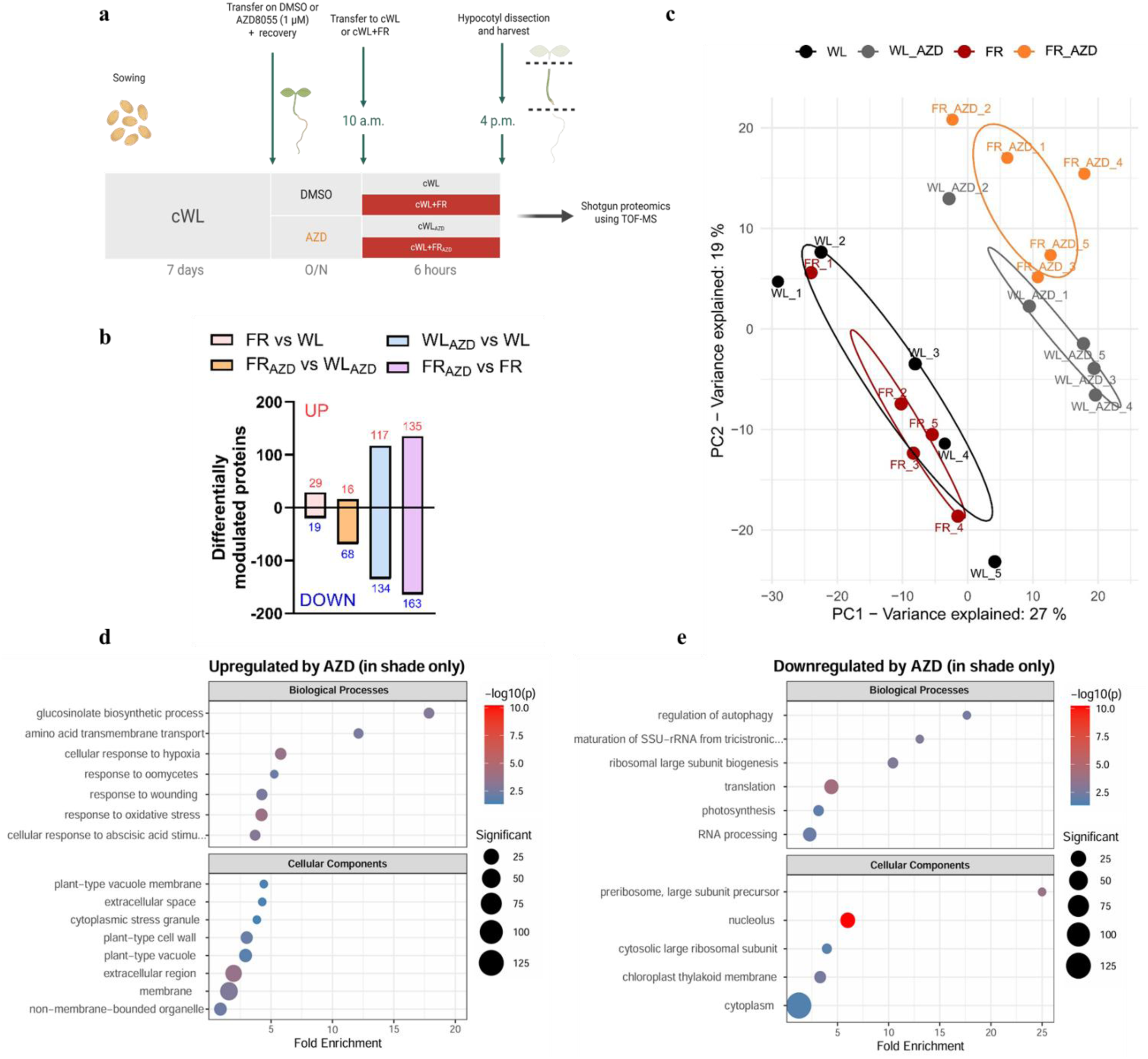
TOR inhibition upregulates stress-related processes and downregulates processes associated with translation. **(a)** Scheme representing the experimental set up used for the proteomic analysis. Seeds were sown onto ½ MS plates, stratified and grown vertically for 7 days in continuous white light (cWL) prior to being transferred onto fresh ½ MS plates containing either DMSO (control) or 1 μM AZD8055. Plates were left to recover from the transfer overnight and were exposed to either cWL or cWL supplemented with far-red light (cWL+FR) to simulate shading. After 6h, hypocotyls were dissected and harvested in liquid nitrogen and used for the proteomic study. **(b)** Proteins found significantly upregulated (UP) or depleted (DOWN) according to the comparisons indicated in the legend. **(c)** Principal component analysis (PCA) was performed on the full list of identified proteins including 5 replicates under cWL in the absence and presence of AZD8055 (WL in black, WL+AZD in grey) and under cWL+FR in the absence and presence of AZD8055 (FR in red and FR+AZD in orange). Missing values were replaced by two-step imputation prior to PCA. **(d and e)** Gene ontology (GO) enrichment analysis of differentially **(d)** upregulated and **(e)** downregulated proteins upon TOR inhibition compared with mock in +FR conditions (FR_AZD vs FR). Shades of blue and red indicate the level of significance based on log10 p.value; circle diameter depicts the number of proteins associated to each GO category. Fold change cut-off was set at FC ≥ 0.7. Diagram shown in panel (a) was created using BioRender.

In control seedlings (not treated with AZD8055), 6 h of cWL+FR exposure resulted in the accumulation of 29 and depletion of 19 proteins, with 5 additional proteins exclusively identified in cWL+FR and 7 exclusively found in cWL (FR vs WL, dataset S1; Fig. 5b; Fig S6a). The auxin-responsive proteins AUX1 and WES1 (AT4G14560 and AT4G27260), as well as several major cell wall-modifying proteins, including xyloglucan endotransglucosylase/hydrolases (XTH8, AT1G11545; XTH16, AT3G23730; XTH18, AT4G30280 and XTH22, AT5G57560) accumulated under cWL+FR compare to cWL. Additionally, an XTH previously reported to promotes elongation in shade was exclusively identified under cWL+FR conditions (XTH17, AT1G65310) (dataset S1) ^38^. However, the number of differentially modulated proteins and a principal component analysis (PCA) indicated that inhibition of TOR had a much stronger impact on the proteome than the light conditions (Fig. 5b and 5c, dataset S3 and S4). Next, we performed gene ontology (GO) enrichment analysis (datasets S1-S4). We found that stress-related biological processes, including “glucosinolate biosynthetic process”, “response to wounding”, “cellular response to hypoxia”, “response to oomycetes” and “response to oxidative stress” (Fig. 5d), were overrepresented in AZD8055-treated compared with mock samples under cWL+FR conditions. In contrast, no stress-related GO categories were found enriched in response to AZD8055 treatment under cWL conditions, except for “cellular response to hypoxia”. These observations suggest that TOR inhibition restores the plant’s capacity to induce stress-related processes, usually suppressed under shade conditions and active TOR. Interestingly, the majority of proteins underrepresented in AZD8055-treated compared with mock samples under cWL+FR, were associated with the GO terms “ribosomal RNA maturation”, “ribosome biogenesis”, “ribosomal subunit precursors” and “translation” (Fig. 5e). Finally, only a few GO categories were associated with proteins differentially enriched under TOR inhibition between cWL+FR- and cWL-treated samples (accumulation of 16 proteins and depletion of 68 proteins; dataset S2). Still, the majority of these GO terms were associated with proteins depleted under shade conditions, including “structural constituent of ribosome”, “cytosolic large ribosomal subunit”, “ribosome” or “translation” (Fig. S6b). Taken together, these results indicate that AZD8055 treatment impairs translation-related processes, particularly under shade conditions.

### TOR inhibition limits shade-mediated hypocotyl elongation via reduced translation efficiency

To explore whether TOR inhibition affects translation efficiency in seedlings grown under shade conditions, we performed polysome profiling of hypocotyls dissected from 7-day-old seedlings exposed for 24 h to either cWL or cWL+FR in the presence or absence of AZD8055 (Fig. 6). The polysome fraction as well as the polysome to monosome (P/M) ratio, indicative of translation efficiency, was strongly enriched in cWL+FR compared with cWL control samples. Importantly, such an increase in heavy polysome fractions was not observed for AZD8055-treated samples, irrespective of light conditions (Fig. 6). Altogether, our data show that shade exposure enhances global translation efficiency in a TOR-dependent manner.

**Figure 6:**
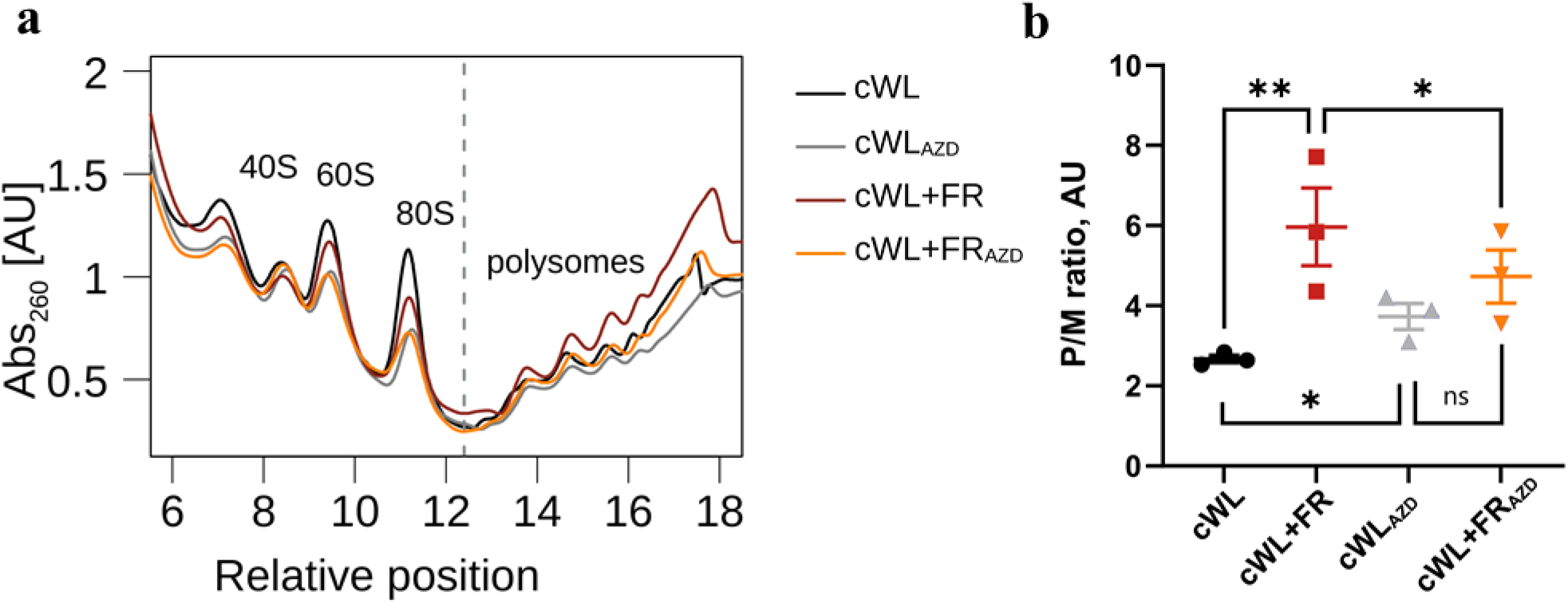
Intact TOR activity is required for shade-mediated activation of global translation. **(a)** Polysome profiling was performed on dissected hypocotyls of 7-day old Col-0 seedlings grown under continuous white light (cWL) or grown under cWL for 6 days followed by 24 h of cWL supplemented with far-red light (cWL+FR), in the presence or absence of the TOR inhibitor AZD8055. The graph is a representative image of three independent experiments. **(b)** Polysome-to-monosome (P/M) ratio calculated from three independent biological replicates. AU, arbitrary units. ns, non-significant. * and ** correspond to *p value* < 0.05 and <0.01, respectively according to two-way ANOVA with Tukey’s multiple comparison test. Graph represents mean values ± SEM (n = 3).

## Discussion

Plant TOR was discovered in the early 2000s ^11^, and since then, a myriad of modes of action and molecular targets have been uncovered, linking TOR to photosynthesis, hormonal balances, translation, energy management and more recently, growth-defense trade-offs ^17,18,39^. In this study, we investigated whether and how TOR contributes to the regulation of shade-induced hypocotyl elongation, using *Arabidopsis thaliana* (Arabidopsis) seedlings as a model. We demonstrate that TOR plays an essential role in shade-induced elongation and define processes affected by both shade and TOR that modulate hypocotyl growth.

Previous work in tomato has shown that *TOR* transcript levels are upregulated shortly after shade exposure ^31^. In this study in Arabidopsis, we found that TOR activity was strongly enhanced by shade compared with white light control conditions, correlating with the activation of shade-induced elongation growth (Fig. 1). We further highlight the need of TOR activation for proper shade elongation responses, as genetic and chemical TOR inhibition leads to defects in elongation growth (Fig. 2). Given the limited shade-induced elongation response in TOR-inhibited plants, we initially hypothesized that TOR inhibition dampens shade-induced auxin responses. However, we were surprised to find that *pDR5::GUS* activity, used as a proxy for auxin-mediated transcriptional responses, was increased in hypocotyls upon shade treatment combined with TOR inhibition, compared with shade alone, showing that initial shade-mediated auxin responses occur independently of TOR (Fig. 4a and 4b). Similarly, when focusing on the hypocotyl region, most shade- and auxin-responsive genes tested showed their highest expression levels under shade combined with TOR inhibition (Fig. 4e-l). Although these results seem counterintuitive, they are consistent with our proteomics data. Indeed, we found that AZD8055 treatments causes depletion of proteins associated to “intracellular auxin homeostasis” including two members of the Gretchen Hagen3 (GH3) family involved in auxin conjugation (GH3.5; AT4G27260 and GH3.6; AT5G54510), suggesting that auxin homeostasis rather than auxin biosynthesis is impaired upon TOR inhibition (dataset S2; Fig. S6b).

Next, we showed that TOR inhibition was associated with the downregulation of processes related to translation and ribosome biogenesis, particularly under shade conditions (Fig. 5d, Fig. S6b). In line with these observations, shade perception was associated with increased translational activity, which is strongly impaired upon TOR inhibition by AZD8055 (Fig. 6). Together, these findings suggest shade-induced TOR activation is required for appropriate hypocotyl elongation under shade conditions. Rapid TOR activation by shade and the subsequent increase in global translation efficiency may allow plants to respond more rapidly and accurately to changing light environments. In contrast, TOR-inhibited plants, which fail to enhance translation efficiency, may be unable to process the mRNA pool fast enough to sustain the growth response. This limitation could lead to increased transcriptional activation as a compensatory mechanism, consistent with the stronger transcriptional response after 24 hours (Fig. 4e-l). Ultimately, this impaired translational capacity may contribute to the growth phenotype observed after 72 hours (Fig. 2a and 2b, see hypothetical model in Fig. 7).

**Figure 7:**
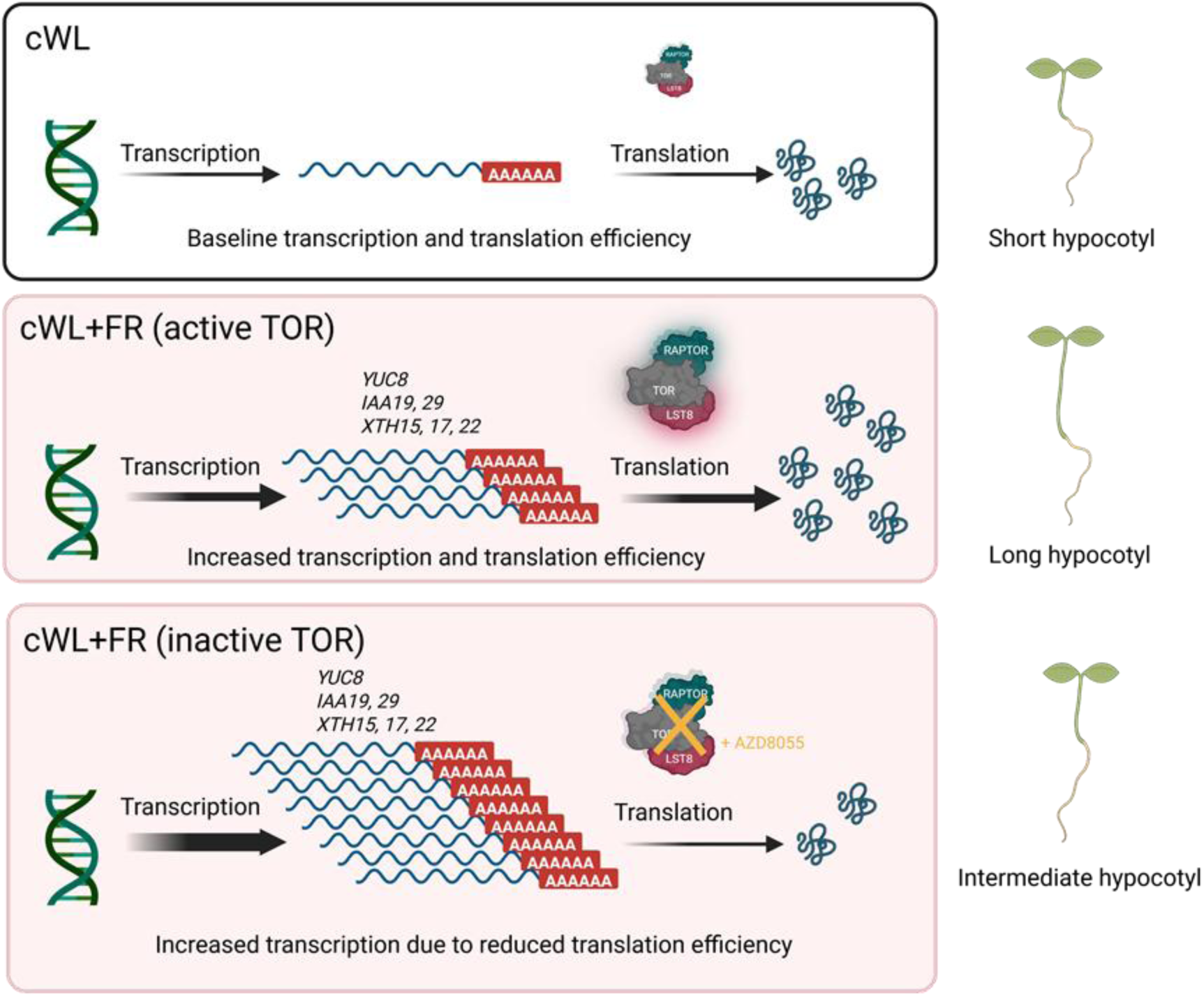
Hypothetical model of how TOR activity modulation by shade impact hypocotyl elongation via differential translation efficiency. In continuous white light (cWL), seedlings display short hypocotyls as a result of basal transcription, translation efficiency and TOR activity (top panel). Under far-red enriched conditions (cWL+FR), TOR activity is promoted and shade- and auxin-responsive genes are upregulated (*i.e. YUC8, IAAs* and *XTHs*). The activation of TOR under these conditions triggers translational reprogramming required for the proper translation of light signaling components and the subsequent hypocotyl elongation response (middle panel). TOR inhibition in shade limits translation efficiency and therefore leads to the accumulation of shade- and auxin-responsive gene transcripts as part of a feedback loop mechanism. TOR-inhibited seedlings hypocotyl elongation capacities in response to shade are thereby reduced (bottom panel). This scheme was created using Biorender.

TOR is known to control ribosome biogenesis, polysome content and translational reprogramming ^40–42^. The latter has been reported in the context of flooding stress tolerance, where general translational is repressed upon submergence, with the exception of specific mRNAs whose translation is increased ^43^. Another study showed monochromatic far-red light, through phyA, also increases the fraction of RNA associated with polysomes ^29^, but other light conditions have not been investigated. We found that shade conditions increase the fraction of polysome-associated RNA in a TOR-dependent manner. To follow up on these observations, translatome profiling (*i.e.* Ribo-Seq, TRAP-Seq) would help unravel the degree of specificity in the shade-TOR relationship. Furthermore, the temporal and spatial impacts of the shade-TOR relationship on elongation growth is still unknown. At the proteome level, we observed only minor differences between shade- and cWL-treated plants after 6 hours (Fig. S6a). This may be explained by the fact that we focused on the hypocotyl, which at this stage may not yet have received all molecular signals and components required for growth originating from the cotyledons, or because these signals remain too localized (*i.e.* in the epidermal cells).

Upon shade perception, auxin is synthesized in the cotyledons and transported to the elongating organs via plasma membrane-localized PIN auxin efflux carriers. Among these, PIN3 is the most extensively studied in the context of shade-induced hypocotyl elongation and petiole hyponasty movement ^4,6,7,44^. While we did not find any PINs among the proteins differentially regulated by shade or TOR inhibition in our proteomics dataset, these treatments may still affect posttranslational modifications of PINs or change their subcellular localization. In roots, glucose-activated TOR has been shown to interact with and phosphorylate PIN2 to control auxin distribution and cell elongation ^45^. It is tempting to hypothesize that other PINs are regulated by a similar mechanism, including PIN3 in the hypocotyl, where TOR may be essential for PIN3 relocalization upon shade perception, allowing auxin transport from the vasculature to the epidermis and promoting elongation ^4,5^. Our data indicate that auxin signaling is misregulated upon TOR inhibition, especially in shade (Fig. 4). While shade acts upstream of auxin and TOR is generally considered a downstream component of auxin signaling, our findings suggest that these pathways may interact within a multilayered regulatory network to fine-tunes shade growth responses. (Fig. 2; Fig. 4).

TOR activity has recently been proposed to be important for elongation growth through the acid-growth mechanism. Active TOR is thought to facilitate water movements into cells, contribute to the induction of cell wall remodeling components, including XTHs, and differentially activate the H^+^-ATPase, leading to apoplastic pH acidification ^46^. Consistent with these ideas, treatment with Fusicoccin only partially rescued the effect of TOR inhibition on elongation growth (Fig. 3e), indicating that TOR activity dynamics during shade perception are linked to hypocotyl epidermal cell elongation, either indirectly or via multiple pathways. Altogether, these observations point to a major role for TOR in cell elongation and possibly cell-to-cell communication, thereby leading to proper hypocotyl elongation under shade conditions ^14^.

Both shaded plants and phytochrome mutants are known to accumulate soluble sugars ^47–49^, which can also contribute to the modulation of TOR activity ^24^. Sucrose-activated TOR has recently been shown to alleviate shade-induced inhibition of leaf development in a phyA-dependent manner ^30^. Interestingly, this study reported a decrease in TOR activity under deep shade conditions (low PAR combined with low R:FR), in contrast to the increased TOR activity in our study under moderate shade conditions (PAR unchanged combined with low R:FR). This discrepancy is likely due to reduced photosynthesis and photosynthate levels under low PAR conditions. Understanding the relationship between TOR and sugar homeostasis and reallocations upon shade perception is therefore key to identifying new molecular targets or processes activated by shade and TOR and contributing to elongation growth. Also, the antagonistic relationship between TOR and SnRK1, activated by energy and nutrient depletion, requires further investigation.

Active TOR not only plays a role in promoting growth but has also been associated with reduced defense responses in different pathosystems ^50–53^. This dual regulation potentially place TOR at the nexus of growth-defense trade-offs to enable rapid and finetuned decision-making in changing environments. We therefore propose that TOR activation by shade may play a role in the reduced defense capacity observed in shaded plants, a phenomenon referred to as the “shade-induced susceptibility” ^48,54–56^. However, further efforts are necessary to pinpoint the molecular hubs associated with TOR that control plant decisions to either grow or defend.

## Conclusion

To conclude, our study reveals a strong link between shade perception and TOR activity modulation in plants. We demonstrate that active TOR is required to trigger proper hypocotyl elongation responses under shade, likely through increased translational efficiency. While the regulation of plant environmental plasticity by TOR relies on a complex and multilayered regulatory network, we managed to bridge multiple research fields to shed light on how TOR activity dynamics impact adaptive growth responses in plants. Finally, we also found that TOR activity is enhanced in shade-exposed tomato plants (Fig. S4) suggesting that its role in shade-induced growth is conserved. Furthermore, our data show that TOR is also involved in growth under elevated temperature. Therefore, targeting TOR in breeding programs could contribute to the development of crop plants better adapted to climate change and high-density planting.

## Materials and methods

### Plant material

Arabidopsis seeds of Col-0 (wild type), *raptor78* (SALK_078159) ^57^, as well as the estradiol inducible TOR RNAi *tor-es1* (N69830) ^34^, *pDR5::GUS* (kindly provided by Dr. Stefan Kircher; Freiburg University, Germany) were used throughout the study.

### Plant growth and light conditions

Arabidopsis seeds were surface sterilized with ethanol and sown onto half strength Murashige & Skoog (½ MS) medium using a sterile toothpick, stratified for >2 days at 4 °C in darkness and grown vertically in HettCube 400R incubators (Hettich) at 21 °C in continuous white light (cWL, 35 μmol.m^-2^.s^-1^) for 4 days to ensure synchronized germination and homogeneous seedling development. Four-day old seedlings were transferred onto fresh ½ MS plates supplemented with either DMSO as a mock treatment, the TOR inhibitor AZD8055 (Hycultec), Kakeimide (ProbeChem), or the synthetic auxins NAA (Sigma) or picloram (MedChemExpress). Concentrations are indicated in figure captions. Unless stated otherwise, the concentration of AZD8055 was kept at 1 μM. Plates were then kept in either cWL (control) or transferred to cWL supplemented with 30 μmol.m^-2^.s^-1^ of FR light (cWL+FR) to simulate upcoming competition. Light spectra and growth procedure are shown in Fig. S1 and S3, respectively.

For experiments with adult plants, *Arabidopsis thaliana* Col-0 seeds were directly sown on wet soil, stratified for >2 days at 4 °C in darkness, and transferred to a growth chamber at 21 °C with white light (WL ∼100 μmol.m^-2^.s^-1^) and long-day photoperiod (16/8 h) to promote germination and accelerate early seedling development. Ten-day-old seedlings were transplanted separately into single round pots (Ø 6 cm) and further grown in short-day conditions (8/16 h) for 4 weeks under white light (WL ∼100 μmol.m^-2^.s^-1^). Four-week-old plants were acclimated for at least 24 h in HettCube 400R incubators (WL ∼120 μmol.m^-2^.s^-1^). At ZT=2 (11 a.m.), half of the plants was exposed to WL supplemented with far-red light (730 nm; 50 μmol.m^-2^.s^-1^), while the other half remained under cWL. Sampling for immunoblotting was performed by harvesting a young developed leaf right before (0 h) and 6 h after the light treatment started. The same leaf was used for petiole elongation using a digital caliper, and for leaf angle measurements after 24 h based on images taken from the side.

*Solanum lycopersicum c.v.* Moneymaker seeds were sown onto wet vermiculite substrate, stratified for >2 days at 4 °C in darkness and transferred to a growth chamber at 21 °C with white light (WL ∼130 μmol.m^-2^.s^-1^) and long-day photoperiod (16/8 h). Seven-day-old seedlings were transplanted into separate square pots and grown for 4 weeks in the same light conditions. Shade treatment were achieved by placing the plants in large open-top white plexiglass boxes equipped with far-red LEDs (FR) mounted on top, allowing top FR illumination while still permitting ambient white light to penetrate. Control plants were placed in the same boxes without the far-red LEDs on. FR supplementation treatment started at ZT=3 (10 a.m.). Measurements were performed on the petiole of the fifth leaf from the bottom, the first internode (right above the hypocotyl) as well as the whole stem.

### Protein isolation and immunoblotting

Samples were harvested in liquid N_2_ and ground using an Ivoclar Silamat S6 for 3×10 sec with freezing in liquid N_2_ in between. Preheated (95 °C) extraction buffer (65 mM Tris-HCl, pH=6.8; 4 M Urea; 10 % Glycerol; 3 % SDS; 0.05 % Bromophenol blue, 10 mM dithiothreitol) supplemented with 10 µl.ml^-1^ protease inhibitor cocktail (Sigma Cat. No P8340), 5 mM sodium fluoride, 2.5 mM sodium pyrophosphate and 5 mM sodium orthovanadate to prevent protein degradation and dephosphorylation. The hot extraction buffer was directly added onto frozen samples (2.5 µl of extraction buffer per 1 mg of sample fresh weight), and samples were vortexed vigorously until homogeneous. Lysates were then incubated under constant shaking on a heat block for 5 min (95 °C, 400 rpm) and centrifuged at 13700 rpm for at least 15 min. The supernatant (protein extracts) was collected in a fresh tube and stored at −80 °C until further analysis.

For protein quantification, 10 μl of extract were added to 190 μl ddH2O and 1 ml of amidoblack staining solution (90 % Methanol, 10 % acetic acid, 0.05 % amidoblack (w/v)). The samples were centrifuged at 13700 rpm, and the pellets were rinsed with washing buffer (90% ethanol, 10% acetic acid), air-dried and dissolved in 0.2 M NaOH. The absorbance at 595 nm was measured using a plate reader. Protein concentrations were estimated based on a Bovine serum albumin (BSA) standard curve.

For immunoblotting, 10-25 μg of total protein extracts were separated on a 10% SDS-PAGE gel and blotted onto polyvinylidene fluoride membranes. Blocking of membranes was performed using 5% BSA in TBS buffer (50 mM Tris/HCL pH= 7.5, 500nM NaCl) supplemented with 0.05 % Tween 20 (TBS-T). Membranes were shortly rinsed with TBS-T prior to overnight incubation with either polyclonal α-RPS6A primary antibody (40S ribosomal protein S6-1) or polyclonal α-RPS6A-P240 primary antibody (phosphorylated (Ser240) 40S ribosomal protein S6-1) from Agrisera (AS19 4292 and AS19 4302, respectively), developed according to previous results ^32^ diluted 1:1000 in TBS-T. The next morning, membranes were washed 3 x 10 min with TBS-T and incubated for one hour at room temperature with α-rabbit peroxidase secondary antibody (Sigma, Cat No A0545) diluted 1:25000 in TBS-T. Membranes were washed again 3 x 10 min with TBS-T, then incubated in TBS until imaging. Imaging was performed using BioRad ECL SuperBright chemiluminescent detection reagents (Agrisera, CAS no AS16 ECL-S). Amidoblack staining was used as a loading control. Signal intensities of bands were quantified using Fiji. A rectangle was drawn around the first band of interest (Analyze > Gels > Select first lane), and then moved to the next bands (Analyze > Gels > Select next lane), before plotting each lane to measure the area under the curves using the wand tool (Analyze > Gels > Plot lanes). The area was taken as the signal intensity for each band and normalized to the intensity of the cWL control. Fold-change in signal intensities compared with cWL was obtained within each blot (α-RPS6 and α-RPS6-P240, respectively). For each condition, the ratios obtained for α-RPS6-P240 were normalized to the ratios obtained for α-RPS6 as indicated below:

Comparison of signal intensity for RPS6 and p-RPS6 separately (normalized value _cWL_ = 1.00):

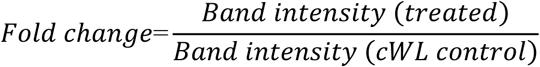

Ratio p-RPS6/RPS6 (ratio is set to 1 in cWL):

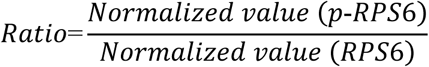

### Gene expression analysis by qPCR

Total mRNA was isolated from ground frozen tissue using an ISOLATE II RNA Plant Kit (Meridian Bioscience) and concentrations were measured using a NanoDrop 1000 spectrophotometer (ThermoScientific). One μg RNA was reverse transcribed into cDNA using a High-Capacity cDNA Reverse Transcription Kit (ThermoFisher Scientific) according to manufacturer’s guidelines. Gene expression analysis was performed by qPCR using qPCRBIO SYBGreen Blue Mix (PB20.17-51 PCR Biosystems) from 10 ng of cDNA per sample using a CFX384 qPCR cycler system (BioRad) with an initial denaturation step (95 °C for 5 min), followed by 40 cycles (95 °C for 30 sec, 60 °C for 1 min, 72 °C for 30 sec). Actin was used as house-keeping gene. Primer sequences are shown in dataset S5.

### Histochemical localization of GUS reporter activity

Seeds from *pDR5::GUS* (in Col-0 background) were grown as indicated previously with minor modifications. Seedlings grown for 7 days on ½ MS plates in cWL and freshly transferred to ½ MS plates supplemented with DMSO or AZD8055 were left in cWL overnight to recover and ensure TOR inhibition prior to starting the light treatment. The next morning, plates were either kept in cWL (35 μmol.m^-2^.s^-1^) or exposed to cWL+FR (35 μmol.m^-2^.s^-1^ WL + 30 μmol.m^-2^.s^-1^ FR) for 24 h. The seedlings were then fixed in ice-cold 90% acetone for 20 min with gentle agitation, and incubated 2x 10 min in washing buffer 0.1 M Phosphate buffer (0.1 M NaH_2_PO_4_, 0.1 M Na_2_HPO_4,_ pH=7.0) mixed with 10 mM EDTA, 2 mM K_3_Fe(CN)_6_) under vacuum, prior to incubation in staining solution (0.1 M Phosphate buffer, 10 mM EDTA, 1 mM K_3_Fe(CN)_6_, 1 mM K_4_Fe(CN)_6_*3H_2_O and 0.5 mg.ml^-1^ X-Gluc) for 10 min under vacuum, followed by overnight incubation at 37 °C. Samples were kept in darkness throughout the procedure. The staining was stopped by adding acetic acid/ethanol (3:1 v/v) for 60 min. The samples were rehydrated by consecutive 5-10 min washes in 75%, 50%, 25% ethanol and 5% ethanol/25% glycerol, respectively. Seedlings were mounted in 50% glycerol on microscopy slides, sealed with transparent nail polish, and incubated for at least 1 hour at 60 °C before observation at the microscope. Images were acquired using a Nikon Eclipse Ni-E light microscope (10x magnification). Image analysis and signal quantification were performed using ImageJ. Color channels were split (Image > Color > split channels) and quantification was performed on the red channel only to have better specificity and sharpness of the GUS signal. Mean and max grey values were obtained from a 250 px wide line drawn across the hypocotyl (Edit > Selection > Line to Area > Measure).

### Protein isolation and quantitative proteomics

Col-0 seeds were sown onto square plates containing ½ MS and stratified for 2 days at 4 °C in the dark. Seedlings were grown vertically on plastic racks in continuous WL (35 µmol m^-2^ s^-1^) for 7 days. Next, seedlings were transferred onto fresh plates containing ½ MS supplemented with either 1 μM AZD8055 or DMSO as a control. In order to guarantee TOR inhibition prior to the shade treatment, seedlings were left to recover from the transfer overnight. The next morning the plates were exposed to either cWL or cWL supplemented with 30 µmol m^-2^ s^-1^ of far-red light to simulate upcoming competition (cWL+FR). The hypocotyls were dissected (roots and cotyledons removed) and harvested in liquid nitrogen. Samples were immediately stored at −80 °C until protein extraction.

Proteins were extracted in SDS-based buffer (4% SDS, 50 mM Tris-HCl, pH6.8, 2.5 mM EDTA, 1% Plant protease inhibitor cocktail (Sigma P9599)) and quantified using the Pierce^TM^ BCA Protein Assay Kit (ThermoScientific). All lysates were adjusted to a concentration of 0.5 mg/ml. 10 µg of each sample were denatured, reduced with 10 mM DTT at 70°C for 10 min, alkylated with 50 mM chloroacetamide at room temperature (RT) for 30 min and quenched with 50 mM DTT for 20 min at RT. Proteome samples were transferred to a 96-deep-well plate (Storage plate 96 well, 1 mL, Agilent Technologies) and desalted with a SP3 bead-based purification protocol ^58^ and digested with trypsin using a STARlet automated liquid handling system (Hamilton) equipped with a 96 well magnet plate (Magnum Flex, Alpaqua). Proteins were bound to Sera-Mag SpeedBead magnetic carboxylate-modified particles (Cytiva) by incubation with 100 µg beads in 80% ethanol for 20 min at RT. After binding, samples were washed three times with 80% ethanol. Proteins were digested overnight at 37°C in 50 µL of 100 mM ammonium bicarbonate containing 0.15 µg of trypsin (Promega V5111). Digestion was stopped the following day by adding 30 µL of 5% formic acid (FA). Peptides were desalted using self-packed SDB-RPS Stop and Go Extraction tips (StageTips; ^59^) composed of three layers of 1.0 × 1.0 mm (AttractSPE® Disks Bio RPS, AFFINISEP). The StageTips were equilibrated successively with 100% methanol, 80% acetonitrile (ACN) containing 0.1% FA, and twice with 0.1% FA. Acidified peptide solutions were applied to the StageTips and washed with 0.1% FA, 80% ACN with 0.1% FA, and 80% methanol with 0.5% FA. Peptides were eluted using 5% ammonia in 60% ACN. The eluates were dried using a SpeedVac concentrator (Eppendorf), reconstituted in 0.1% FA and diluted 1:5 in 0.1% FA.

### Mass spectrometry data acquisition

Samples were separated on a Vanquish UHPLC (Thermo) equipped with a PepMap Neo trap cartridge (0.5 cm, 300 µm inner diameter, C18, Thermo) and a µPAC Neo chromatography column (50 cm, 75 µm inner diameter, C18, pore size <100 Å, Thermo) and directly analyzed with a timsTOF Ultra2 mass spectrometer (Bruker) coupled to chromatography system with a CaptiveSpray 2 ion source (Bruker) equipped with a 20 µm emitter. Peptides were loaded onto the trapping column at a flowrate of 20 µL/min in 3% solution B (80% ACN, 0.1% FA). The trapping column was connected to the separation column at a flowrate of 500 nl/min and after 1 min the peptides were eluted over 41 min at a flowrate of 300 nl/min in a linear gradient starting from 8% solution B ending at 42% B. Subsequently, solution B was increased to 99% within 12.5 min. The timsTOF Ultra2 was operated in DIA-PASEF mode with a capillary voltage of 1600 V, an MS1 scan range of 100 to 1700 m/z and a tims range of 0.64 to 1.45 V*s/m^2^ (1/k0). Ramp and accumulation time were set to 100 ms at a ramp rate of 9.34 Hz. One MS cycle contained 1 MS1 and 8 MS/MS ramps and a total of 24 MS/MS windows covering an m/z range of 400-1000 at and a tims range from 0.64 to 1.37 V*s/m^2^.

### Mass spectrometry data analysis

Raw data were searched using DIANN 2.3.0 ^60^ against a predicted library containing all Araport11 protein models ^61^ as well as sequences of known contaminants. Settings for library generation and database search were a peptide length from 7-35 amino acids within an m/z range of 390 to 1010 and a precursor charge range from +1 to +4. The fragment mass range was set from 150 to 1600 m/z. The protease specificity was set to trypsin (C-terminal K|R) with a maximum of 1 missed cleavage. Carbamidomethylation of cysteines (+57.021464 Da) was specified as fixed modification and N-terminal acetylation (protein N-terminus, +42.010565 Da), oxidation of methionine (+15.994915 Da) and deamidation of asparagine (+0.984016 Da) were specified as variable modifications. A maximum number of 2 variable modifications was allowed. For the database search, the MS1 accuracy was set to 10 ppm and the MS2 accuracy was set to 12 ppm. Quantitative analysis using the protein and peptide matrices from DIANN was performed in RStudio ^62^. First, the dataset was restricted to proteins identified in at least three of five replicates in each condition of the performed comparison. LFQ intensities were log2-transformed and analyzed for differential abundance using the limma package ^63^ with empirical Bayes moderation and Benjamini-Hochberg correction for multiple testing. GO term enrichment was performed using topGO ^64^ on proteins whose abundance was significantly changed in a condition including proteins exclusive to the respective condition (at least 3 values in the condition and zero values in the compared condition, see tables “exclusive proteins in dataset S1).

Principal component analysis (PCA) was performed on the full list of identified proteins. Missing values were handled using a two-step imputation approach: proteins missing in all or more than three replicates of a category were imputed using left-censored minimum probability (MinProb), while proteins missing in three or fewer replicates were imputed using impSeqRob as implemented in the rrcovNA package v0.5.3 ^65,66^.

### Polysome profile analysis

Two hundred hypocotyls per condition were snap-frozen in liquid nitrogen and ground to a fine powder using an Ivoclar Silamat S6 for 3×10 sec with freezing in liquid N_2_ in between. The powder was resuspended in Polysome Extraction Buffer (200 mM Tris-HCl, pH 9.0; 200 mM KCl; 25 mM MgCl_2_; 25 mM EGTA, pH 8.0; 1x Detergent mix [1% Brij-35 (w/v), 1 % Triton X-100 (w/v), 1 % Igepal CA-630 (w/v), 1 % Tween 20 (w/v)], 1 % Polyoxyethylene-10-tridecyl ether (PTE), 1 % Sodium Deoxycholate (DOC), 5 mM dithiothreitol (DTT), 100 µg/mL cycloheximide, cOmplete™ EDTA-free Protease Inhibitor Cocktail [Roche]). Lysates were cleared by centrifugation at 16,000 g for 15 min at 4 °C. The equivalent of 3 µg of total DNA (quantified using Qubit™ DNA High-Sensitivity Assay Kit) was layered on top of the 10 to 60 % (w/v) sucrose density gradients, prepared using BioComp Gradient Master. Ultracentrifugation was carried out at 38,000 rpm for 3 h at 4 °C in a SW41 Ti rotor (Beckman Coulter). Polysome profiles were recorded by measuring the absorbance at 260 nm using a Biocomp Station in combination with Triax software. Profiles were then aligned according to the 80S peak and the polysome-to-monosome (P/M) ratio was calculated using QuaPPro ^67^.

## Quantification and statistical analysis

Statistical analyses were performed using Microsoft Excel, R, and GraphPad Prism. Student’s t-tests were used for datasets containing two samples and were conducted in Microsoft Excel. Analyses involving two or more factors were performed using analysis of variance (ANOVA), followed by Tukey’s post hoc tests, in RStudio (R version 4.3.1) ^68^. Statistical analyses of polysome profiling data were performed using GraphPad Prism 10.

## Data availability

The mass spectrometry proteomics data have been deposited to the ProteomeXchange Consortium (PMID: 36370099) via the PRIDE partner repository (PMID: 39494541) with the dataset identifier PXDxxxxx (Reviewer access token:xxx)

## Declaration of generative AI and AI-assisted technologies

During the preparation of this work, the author(s) used ChatGPT in order to improve the readability and language of the manuscript. After using this tool/service, the author(s) reviewed and edited the content as needed and take full responsibility for the content of the published article.

## Author contributions

SC conceptualized the research, acquired funding, performed the experiments, designed the figures and wrote the manuscript. MS performed the polysome profiling, provided the figures and analysis. SH performed the proteome analysis, handled the raw data and designed the proteomics figures. IL performed the tomato experiments. PH, CM and AH read and provided intellectual support and feedback. All authors read, edited and approved the manuscript prior to submission.

## Acknowledgments

We are grateful to the Department of Molecular Plant Science (DOMPS) community of Freiburg University for their support, intellectual input and feedback on the project. We also wish to thank the plant TOR community as well as the photobiology community for stimulating discussions and useful feedback during the project.

## Supplemental figures and legends

**Figure S1:**
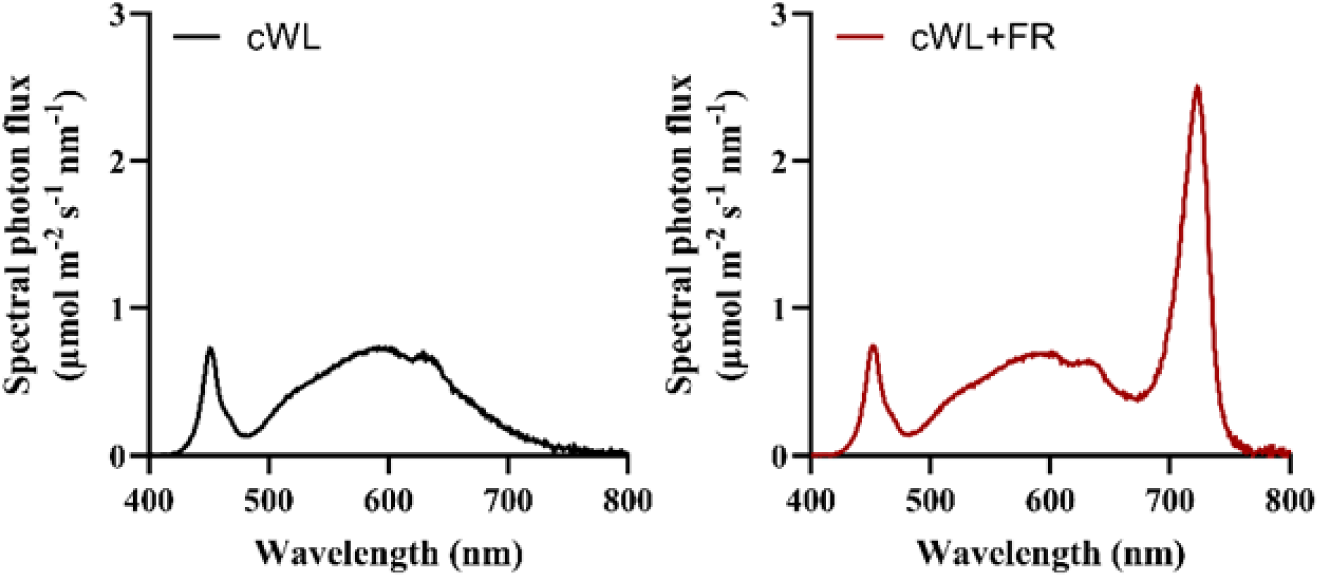
LED spectral composition used throughout the study. Continuous white light (cWL, 35 µmol. m^-2^. s^-1^) was used as a control. Shade treatments were achieved by supplementing a continuous white light background with far-red LEDs (cWL+FR; 30 µmol. m^-2^. s^-1^).

**Figure S2:**
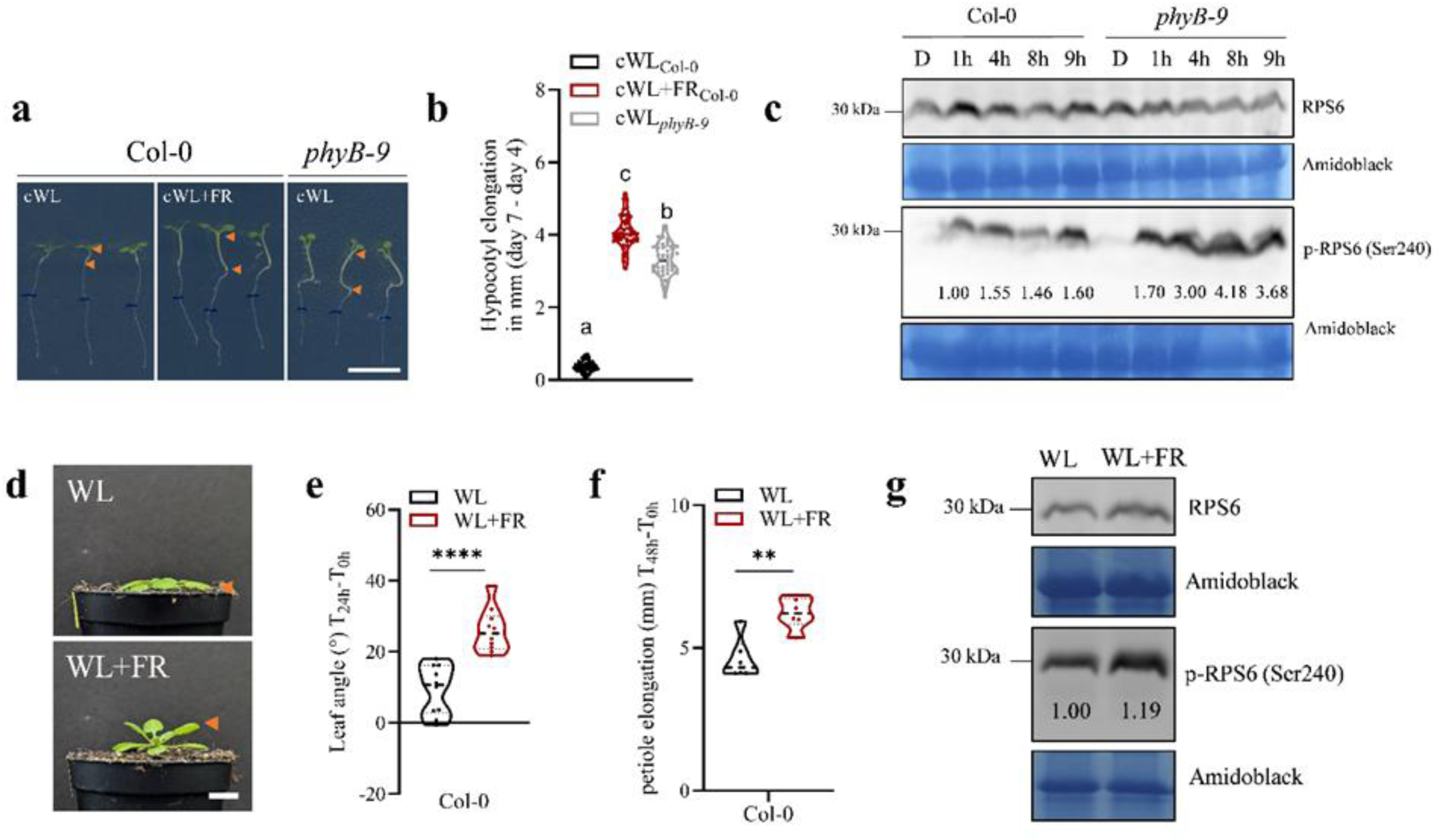
Far-red-mediated TOR activation occurs in adult plants and is linked to phyB signalling. **(a and b)** Representative images and hypocotyl elongation measurement of 7-day-old Col-0 and *phyB-9* mutant seedlings after 7 days continuous white light (cWL), or after 4 days of cWL followed by 3 days in cWL supplemented with far-red LEDs (cWL+FR). n = 45-48 hypocotyls per timepoints and conditions. **(c)** Immunoblotting using polyclonal antibodies targeting RPS6 or a specific phosphorylation site on Serine 240 (p-RPS6 (Ser240)) on crude protein extracts (25 μg) originating from 10-day old Col-0 and *phyB-9* mutant seedlings grown on wet filter paper in short day conditions either in darkness (D) or after 1h, 4h, 8h and 9h of WL. **(d-f)** Representative images, leaf angle and petiole elongation measurement of 3-week-old Col-0 wild-type plants grown in white light (WL; short day; 8h/16h) and treated for 24 h or 48 h with either WL or WL supplemented with far-red light (WL+FR). n = 8-10 plants per conditions. **(g)** Immunoblotting using antibodies against the TOR target 40S ribosomal protein S6 (RPS6A) (RPS6) or against a specific phosphorylation on Ser240 (anti-p-RPS6 (Ser240)) from crude protein extracts (25 μg) originating from Col-0 leaves 6h after the start of the WL or WL+FR treatment. n= 8 plants (1 leaf per plant pooled per sample). Amidoblack staining was used as a loading control. Scale bar = 1 cm. **** and ** corresponds to *p value* < 0.0001 and <0.01, respectively. Letters indicate significant differences according to one-way ANOVA, Tukey’s post-hoc test. Orange arrows indicate leaf of interest (d) and hypocotyls regions (a).

**Figure S3:**
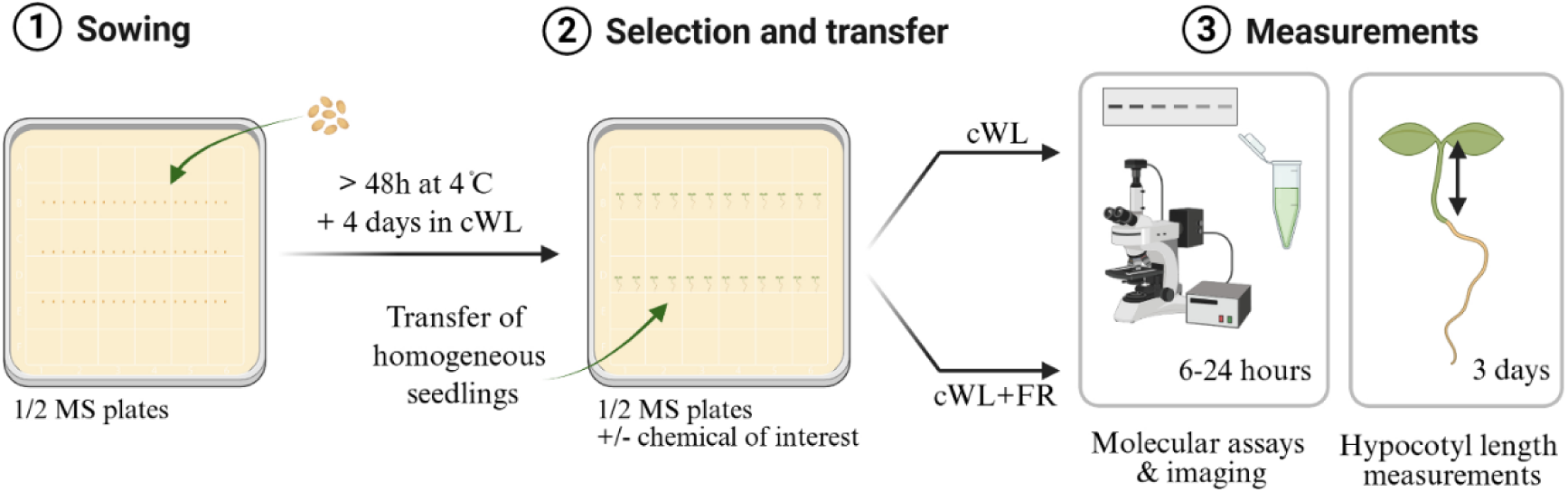
Experimental setup used throughout the study. (1) Seeds were surface sterilized with ethanol prior to sowing onto ½ MS plates (1% agar) using a sterile toothpick. Plates were stratified for at least 48 hours at 4 degrees in darkness before being transferred to continuous white light (cWL) for 4 days to allow for homogenous germination. **(2)** Seedlings were selected and transferred onto fresh ½ MS plates (1% agar) supplemented or not with a chemical of interest (AZD8055, estradiol, etc…). DMSO or ethanol was used as control. Plates were transferred to either cWL or cWL supplemented with far-red LEDs (cWL+FR) immediately after transfer. **(3)** Depending on the experiments, seedlings were harvested at either 6h, 24h (*i.e.* western blots, proteomics, polysome profiling) or 72h (phenotyping). The figure was designed using BioRender.

**Figure S4:**
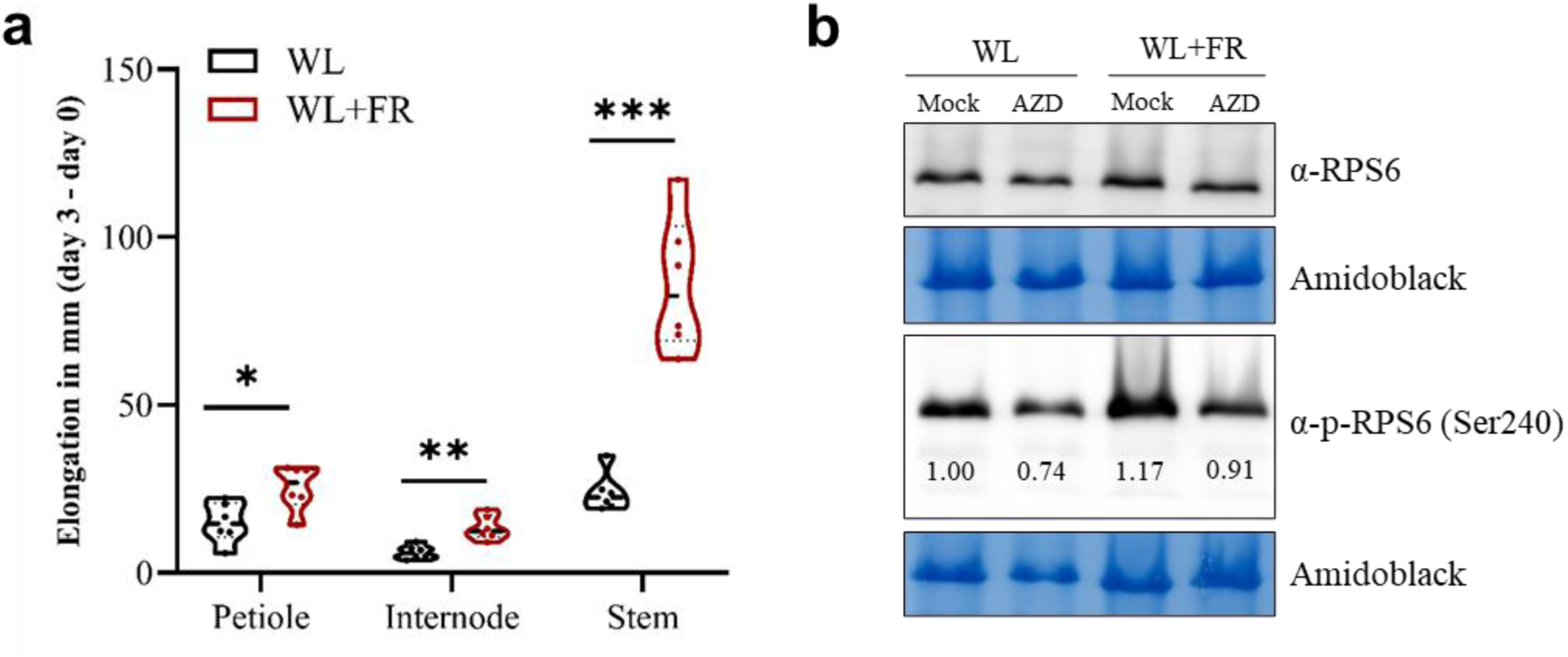
TOR activation by shade is conserved in tomato. **(a)** Relative elongation in 3-week-old tomato *cv.* Moneymaker plants, grown in white light (WL; long day; 16h/8h) and treated for 72 h with either WL or WL supplemented with far-red LEDs (WL+FR). FR supplementation treatment started at ZT=3 (10 a.m.). Measurements were performed on the petiole of the fifth leaf from the bottom, the first internode (right above the hypocotyl) as well as the whole stem. Data show mean ±SEM. n = 6-8 plants per conditions. *, ** and *** corresponds to *p value* < 0.05, <0.01 and <0.001, respectively. Asterisks represent significant difference with Student t.test, p *<0.05, **<0.01, ***<0.001. **(b)** Immunoblotting using polyclonal antibodies against the TOR target 40S ribosomal protein S6 (RPS6A; α-RPS6) and against a specific phosphorylation site at Ser240 (α-p-RPS6 (Ser240)) state from crude protein extracts (25 μg) originating from leaf discs from fifth leaf at 6h after the start of the WL or WL+FR treatment. n= 8 plants (3 leaf discs per plants were pooled per conditions). Amidoblack staining was used as a loading control. Ratio in band intensity between α-p-RPS6 (Ser240) and α-RPS6 was calculated and normalized to the intensity of the WL control for each lane.

**Figure S5:**
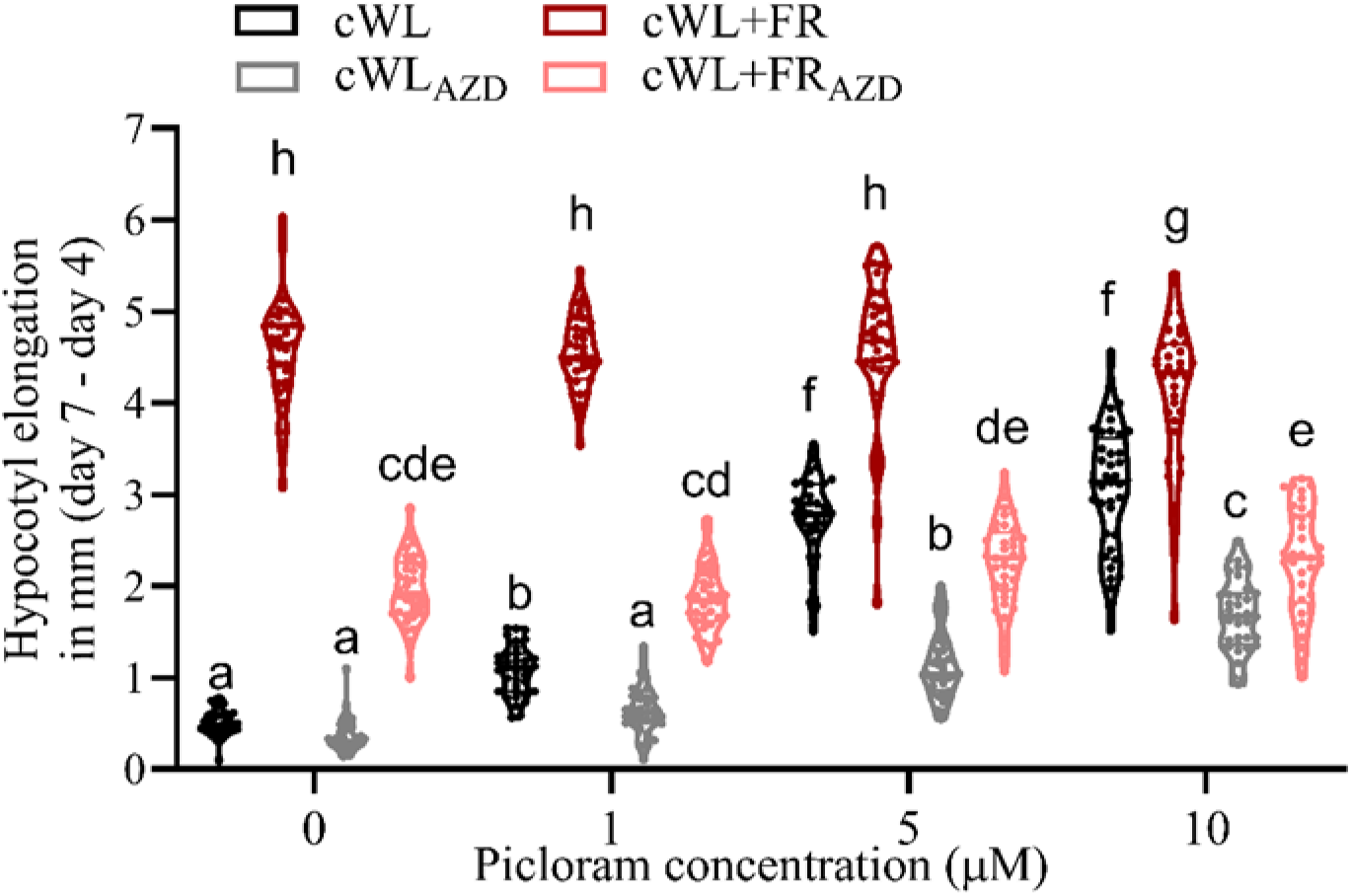
TOR inhibition reduces responsiveness to auxin in Arabidopsis seedlings. Hypocotyl elongation measurement of 7-day-old Col-0 wild-type seedlings after 7 days in continuous white light (cWL), or after 4 days of cWL followed by 3 days in cWL supplemented with far-red light (cWL+FR), and treated with DMSO as a control (0) and with 1, 5 or 10 µM of the auxin analog Picloram. Data represent the elongation between before and after 3 days of cWL or cWL+FR treatment (day 7 – day 4). Letters indicate significant differences according to three-way ANOVA, Tukey’s post-hoc test.

**Figure S6:**
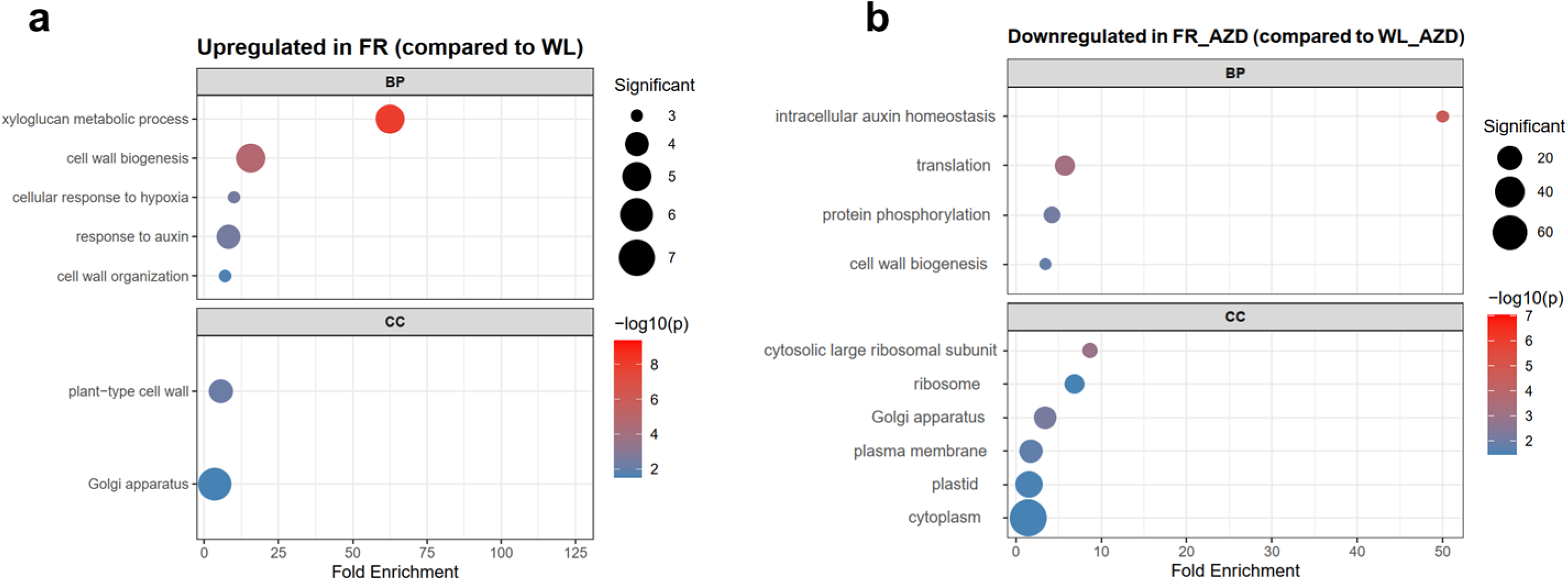
Shade avoidance signatures are present in the hypocotyl after 6 hours and the downregulation of translation by AZD is stronger in FR. Gene ontology (GO) enrichment analysis of differentially **(a)** upregulated proteins upon FR supplementation compared to WL in control conditions and **(b)** downregulated upon FR_AZD treatment compared to WL_AZD. Shades of blue and red indicate the level of significance based on log10 p.value and circle diameter depict the number of proteins associated to each GO category. Fold change cut-off was set at FC ≥ 0.7.

